# NO-stressed *Y. pseudotuberculosis* have decreased cell division rates in the mouse spleen

**DOI:** 10.1101/2021.08.04.455180

**Authors:** Bessie Liu, Rezia Era D. Braza, Katherine L. Cotten, Robert K. Davidson, Kimberly M. Davis

## Abstract

Fluorescence dilution approaches can detect bacterial cell division events, and can detect if there are differential rates of cell division across individual cells within a population. This approach typically involves inducing expression of a fluorescent protein, and then tracking partitioning of fluorescence into daughter cells. However, fluorescence can be diluted very quickly within a rapidly replicating population, such as pathogenic bacterial populations replicating within host tissues. To overcome this limitation, we have generated two revTetR reporter constructs, where either mCherry or yellow fluorescent protein (YFP) is constitutively expressed, and repressed by addition of tetracyclines, resulting in fluorescence dilution within defined timeframes. We show that fluorescent signals are diluted in replicating populations, and that signal accumulates in growth-inhibited populations, including during nitric oxide exposure. Furthermore, we show that tetracyclines can be delivered to the mouse spleen during *Yersinia pseudotuberculosis* infection, and defined a drug concentration that results in even exposure of cells to tetracyclines. We then used this system to visualize bacterial cell division within defined timeframes post-inoculation. We detected growth attenuation of the revTetR-mCherry strains within mouse tissues, however data suggested heightened NO exposure correlated with heightened mCherry signal. We were able to restore normal bacterial growth with revTetR-YFP, and use this strain to show that heightened NO exposure correlated with heightened YFP signal, indicating decreased cell division rates within this subpopulation *in vivo*. This revTetR reporter will provide a critical tool for future studies to identify and isolate slowly replicating bacterial subpopulations from host tissues.

## Introduction

Bacterial growth and metabolism have been the focus of decades of research, in part because we need to learn more about the pathways utilized by bacteria for growth to be able to target these essential processes with antibiotics or novel therapeutics. It is also well appreciated that bacteria with slowed metabolic rates are less susceptible to antibiotics (1–4). Earlier studies focused on growth of bacteria and antibiotic susceptibility under different laboratory conditions (1, 5–7); more recently, there has been a focus on studying bacterial growth and metabolism within host tissues, to more closely approximate bacterial growth conditions at the point where antibiotics or other therapeutics would be administered (8–10). Importantly, bacterial pathogens may utilize a range of different nutrients for growth within host tissues, and individual bacteria, or subsets of bacteria, may concurrently utilize different metabolites to promote their growth (10–14). Some of these substrates may support faster bacterial replication rates, and others, slower replication rates. Nutrient availability and the presence of host-derived antimicrobials, in addition to interactions with different immune cells subsets, can also generate complex microenvironments within host tissues, which can result in differences in growth rates across bacterial populations (12, 15–18).

When treating a bacterial infection, differences in bacterial growth rates can dramatically impact antibiotic efficacy. Slow-growing subsets of bacteria are inherently less susceptible to antibiotic treatment, due to decreased metabolic rates (1, 3, 4). This is termed antibiotic tolerance, a transient, phenotypic change in a subset of bacterial cells within a population, which alters antibiotic susceptibility (19, 20). Many of these experiments have been performed in bacteriological media, so it still remains largely unclear which pathways promote decreased antibiotic susceptibility and slowed growth within host tissues. To identify these pathways, investigators will need to isolate antibiotic tolerant cells from the host environment for downstream analyses. The specific bacterial pathways linked to tolerance may depend in part on the antibiotic and the mode of action, or the environment around the bacteria during the antibiotic exposure (18, 21, 22).

Molecular tools currently exist to identify and isolate slow-growing bacterial cells (8, 23–26), but there are also limitations with some of these approaches that need to be addressed to apply them within the host environment. One of the main approaches is utilizing fluorescence dilution to identify individual cells that have divided within a certain timeframe. Typically, fluorescence dilution approaches involve inducing expression of a fluorescent protein then removing the inducer, and detecting changes in the amount of fluorescent protein within cells as a readout for cell division events and protein partitioning into daughter cells (27, 28). Many research groups have successfully utilized this approach to identify dividing and non-dividing bacterial cells, by inducing fluorescent expression prior to introducing bacteria into tissue culture or infection models (27–30). For some bacterial infection models, this approach can easily be applied, because the size of the colonizing population of bacteria is relatively large, and the number of cell division events within host tissues is relatively small (24, 25, 27). However, in other infection models, only a few bacterial cells establish infection, and these cells go on to replicate many times, which would quickly dilute out fluorescent signals (17, 31–33). Furthermore, it is difficult to deliver inducer compounds into host tissues at sufficient levels to modulate gene expression, and more challenging to remove these compounds to allow for subsequent fluorescence dilution.

*Yersinia pseudotuberculosis* is an enteric pathogen that spreads systemically from intestinal tissues to colonize deep tissue sites, such as the spleen (34–36). There is a strong population bottleneck within intestinal tissues, and few bacterial cells spread systemically to colonize deep tissue sites (37). Within deep tissues, individual bacteria found clonal clusters of replicating extracellular bacteria, termed microcolonies or pyogranulomas (17, 38, 39). Microcolonies can grow to contain hundreds to thousands of bacteria within a single replication site, despite recruitment of circumscribing layers of neutrophils and monocytes (17, 39, 40). Recent studies have shown that subpopulations of bacteria are present within microcolony structures, and that subsets of bacteria stressed by the presence of nitric oxide (NO) preferentially survive treatment with doxycycline, suggesting they may represent a slow-growing subset of cells (17, 22). However, it has been difficult to determine growth rates in this model system, since fluorescence dilution approaches cannot typically be used when the bacterial population contracts and expands so dramatically.

To overcome these limitations, we have generated revTetR reporter constructs, where exposure to tetracyclines will inhibit additional fluorescent gene expression, resulting in fluorescence dilution. These constructs will promote fluorescent signal accumulation in non-dividing cells, and fluorescent protein will be diluted into dividing cells, thus allowing us to differentiate between dividing and non-dividing subpopulations of bacteria. Tetracyclines can also easily be delivered into our mouse model of *Yersinia pseudotuberculosis* systemic infection (22), which allows us to visualize fluorescence dilution as a measure of cell division at specific stages of infection and within specific timeframes. Here, revTetR constructs allow us to ask if responses to host-derived stresses, specifically nitric oxide (NO), impacts *in vivo* cell division rates. These reporter constructs will be a useful tool for future studies to isolate and characterize bacterial subpopulations with differential growth rates.

## Results

### The revTetR reporter can be used to visualize differential bacterial growth rates *in vitro* with mCherry fluorescence dilution

To detect differential cell division rates within host tissues, we generated a fluorescence dilution reporter system by modifying our recently described TetON fluorescent reporter with stable mCherry (*tetR*::*P_tetA_::mCherry*) (22). The revTetR reporter (*revtetR::P_tetA_::mCherry*) was constructed by introducing two mutations (E15A and L17G) into the wild-type *tetR* gene. These two mutations were predicted to cause a reversal of TetR activity resulting in repression of *P_tetA_* specifically in the presence of tetracycline and tetracycline derivatives (41, 42). Therefore, revTetR is a tetracycline-responsive reporter that allows cells to constitutively express mCherry in the absence of tetracycline derivatives, and subsequent addition of tetracyclines should inhibit additional mCherry transcription **(Figure 1A)**. This reporter can be used to compare growth rates of bacteria via a fluorescence dilution approach (27, 28). For example, in cultures treated with tetracycline, fast-growing cells will continue to divide, and the existing mCherry signal will be partitioned between daughter cells with every replication event **(Figure 1B)**. However, slow-growing or nondividing cells will not go through the same number of replication events and therefore will not undergo fluorescence dilution **(Figure 1B)**.

**Figure 1.**
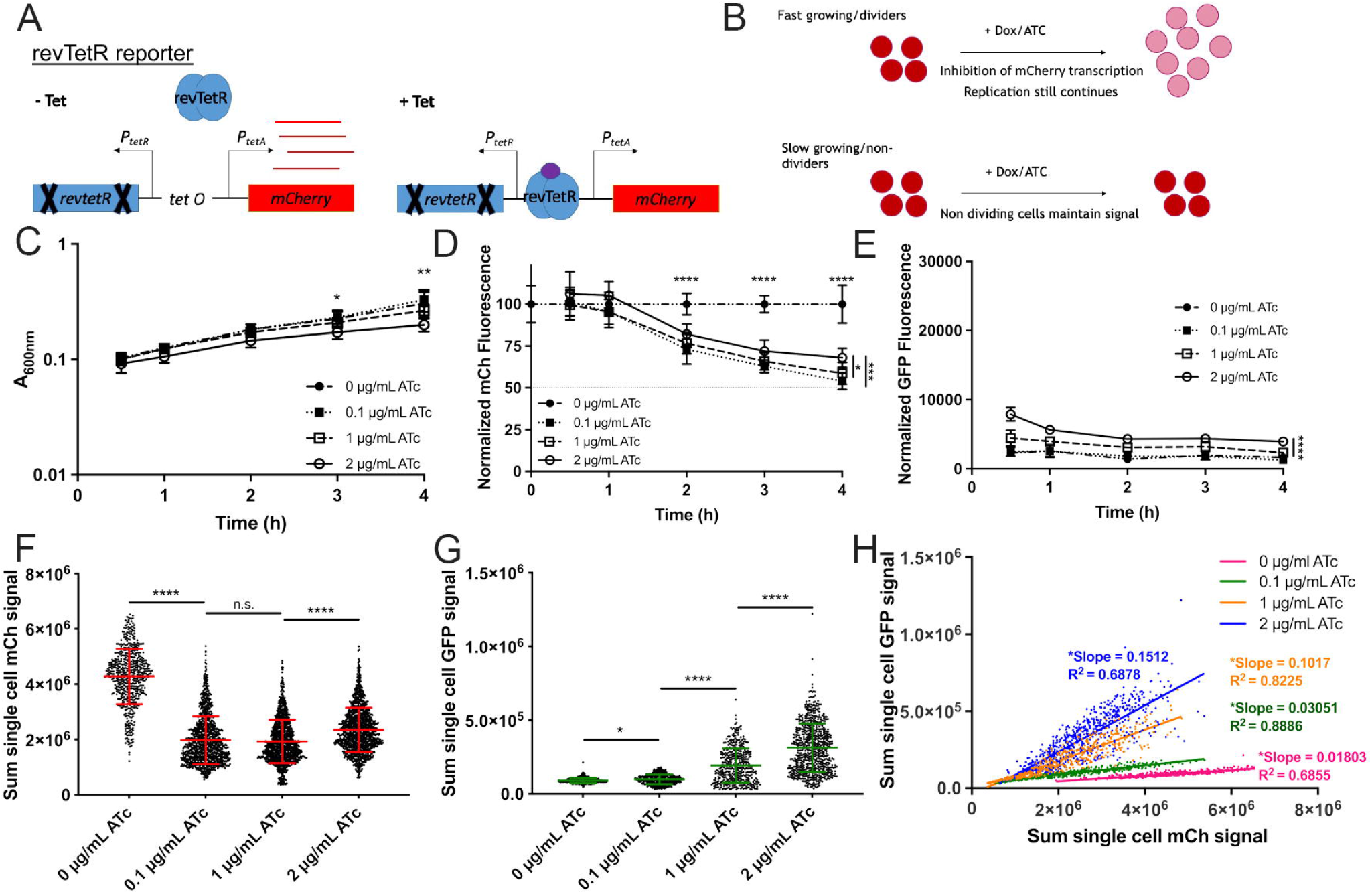
The revTetR reporter can be used to visualize differential bacterial growth rates *in vitro* with mCherry fluorescence dilution. **(A)** The revTetR-mCherry reporter was created by introducing two mutations (E15A and L17G) in the wild-type tetR sequence, which results in constitutive mCherry expression in the absence of tetracyclines (-Tet). Tetracycline addition (+Tet) promotes revTetR binding and repression of mCherry transcription. **(B)** After tetracycline addition (+Dox/ATc), fast-growing/dividing bacteria (top row, red circles) will partition mCherry equally into daughter cells with each cell division event, resulting in dilution. Slow-growing/non-dividing cells will retain a high level of mCherry (bottom row). **(C-E):** Exponential phase cultures of the revTetR-mCherry *dps* strain (2h subculture in LB, 37° C with rotation) were treated with the indicated ATc doses at 0h (hours, h), then further incubated at 37° C with rotation. Aliquots were taken at the indicated timepoints to detect **(C)** optical density by absorbance (A_600nm_), **(D)** mCh fluorescence (normalized by dividing total fluorescence by OD values, untreated value set to 100%), dotted line depicts one doubling (50%), and **(E)** GFP fluorescence normalized to OD. **(C-E)**: Data shown from 9 replicates. **(F)** Cells treated with the indicated doses of ATc for 4h were fixed for fluorescence microscopy. Sum single cell mCherry signal is shown. **(G)** Sum single cell GFP signal is shown. **(H)** Correlation plot depicting single cell sum mCherry and sum GFP signals. **(F-H):** Data shown from three biological replicates. Mean and standard deviation are shown. Statistics: **(C-E)** Two-way ANOVA with Tukey’s multiple comparison test, comparisons made between 2µg/ml and other groups; **(F-G)** Kruskal-Wallis one-way ANOVA with Dunn’s post-test; **(H)** linear regression with R^2^ value and slope of best-fit line, significantly non-zero slope indicates values are correlated. ****p<.0001, **p<.01, *p<.05, n.s.: not-significant.

To confirm that the mCherry fluorescence from the revTetR reporter dilutes into actively dividing cells, we generated a reporter strain containing the revTetR reporter alongside a plasmid containing *dps::gfp-ssrA* (revTetR *dps*) (16). Dps, a ferritin-like iron-sequestering protein, accumulates within bacteria in stationary phase in response to multiple different stresses and protects DNA from oxidative damage (43–45). Slow-growing cells have increased levels of the *dps* (GFP) reporter signal (16). If the revTetR reporter is functioning as predicted, we would expect to detect heightened revTetR (mCherry) in cells expressing *dps,* identifying these cells as slow-growing, non-dividing cells within the population. revTetR *dps* cells were grown into exponential phase then treated with increasing doses of anhydrotetracycline (ATc), a tetracycline derivative that binds TetR or revTetR but has limited ribosomal targeting (46, 47). Aliquots of each sample were measured for optical density or absorbance (A_600nm_) and fluorescence using a microplate reader. The absorbance values increased across all treatment groups, and ATc did not have a detrimental effect upon growth rate with either 0.1µg/ml or 1µg/ml treatment **(Figure 1C)**. However, 2µg/ml ATc treatment did slightly inhibit growth, resulting in significantly lower OD values at 3h and 4h post-treatment (hours, h). mCherry fluorescence values were normalized to cell number by dividing the background-subtracted fluorescence value from each well by the corresponding OD value. The resulting value was further normalized to the untreated sample value for each time point (set at 100%). Following 2h of ATc treatment, all treated groups had significantly reduced mCherry fluorescence relative to untreated cells **(Figure 1D**). At 4h post-treatment, the fluorescent signal for all treated groups decreased by approximately 50% relative to the signal of the untreated group, showing that ATc successfully induces dilution of mCherry with each replication event **(Figure 1D)**. A 50% dilution in fluorescent signal should correlate with one cell division event in all the cells within the population, suggesting the doubling time was approximately 4h. This is slow for *Y. pseudotuberculosis* strains, and was the first indication that the high levels of mCherry expression associated with this revTetR construct may impact bacterial growth. We noted that 2µg/ml ATc treatment resulted in significantly higher mCherry **(Figure 1D)** and *dps* reporter signal **(Figure 1E)** relative to 0.1µg/ml and 1µg/ml treatment, again suggesting the 2µg/ml dose may slightly inhibit growth, and providing evidence that slowed growth can be detected by an increase in mCherry signal. Cells from the final time point (4h) were fixed and imaged using fluorescence microscopy to quantify single cell fluorescence using the sum single cell signal for each channel. The average signal of the untreated group (0µg/ml) was significantly higher than the average signal of the group treated with the lowest dose of ATc (0.1µg/ml), again showing that signal dilution was successful **(Figure 1F)**. 1µg/ml ATc treated cells had mCherry fluorescence similar to the 0.1µg/ml treated group. However, at the highest dose of ATc (2µg/ml), the average mCherry signal signficantly increased **(Figure 1F)**, and we found there were also significant increases in *dps* reporter expression with increasing doses of ATc **(Figure 1G)**, suggesting that high concentrations of ATc may have an inhibitory effect on bacterial cells. Slower growing cells with heightened *dps* reporter expression should also exhibit heightened revTetR-mCherry signal, and we confirmed these signals were correlated under all treatment conditions, based on significantly non-zero slopes of linear regression best-fit lines **(Figure 1H)**. Collectively these data indicate that revTetR fluorescence is accumulating in slow-growing cells, and that actively growing cells are partitioning mCherry into daughter cells, resulting in significant decreases in mCherry signal.

### Growth inhibition with doxycycline promotes mCherry fluorescence accumulation

To further test whether growth-inhibited cells exhibit heightened revTetR-mCherry fluorescence, cells of the revTetR *dps* strain were sub-cultured to exponential phase and treated with increasing doses of the antibiotic, doxycycline (Dox). Aliquots of each sample were measured for optical density (OD) by absorbance (A_600nm_) and fluorescence at hourly timepoints using a microplate reader. Cells exposed to 1µg/mL Dox had significantly decreased growth rates, however the doses less than 1µg/ml (0.01-0.1µg/ml) were subinhibitory, and did not impact growth **(Figure 2A)**. mCherry fluorescence was initially heightened with 1µg/ml Dox treatment **(Figure 2B)**, and *dps* reporter signal was slightly, but not significantly, elevated **(Figure 2C)**, consistent with growth inhibition at this treatment concentration. Three hours after treatment, the mCherry signal in all treatment groups was significantly below untreated values, including the 1µg/ml treatment group **(Figure 2B)**, suggesting that growth may resume after initial growth inhibition. Single cell analysis was performed by fluorescence microscopy, and the sum single cell fluorescence values were graphed for each treatment group. Dilution of mCherry signal could be seen with treatment of 0.01µg/mL Dox, but as the dosage increased, the mCherry signal within cells increased, indicating signal accumulation and stalled growth **(Figure 2D)**. The mCherry fluorescent signal was significantly higher in cells treated with 1µg/ml Dox compared to untreated cells (0µg/ml), suggesting that growth inhibition resulted in significant revTetR-mCherry fluorescence accumulation. Again, we observed a correlation between *dps* reporter signal and revTetR-mCherry signal under all treatment conditions, confirming that slower growing cells have heightened revTetR fluorescence **(Figure 2E)**.

**Figure 2.**
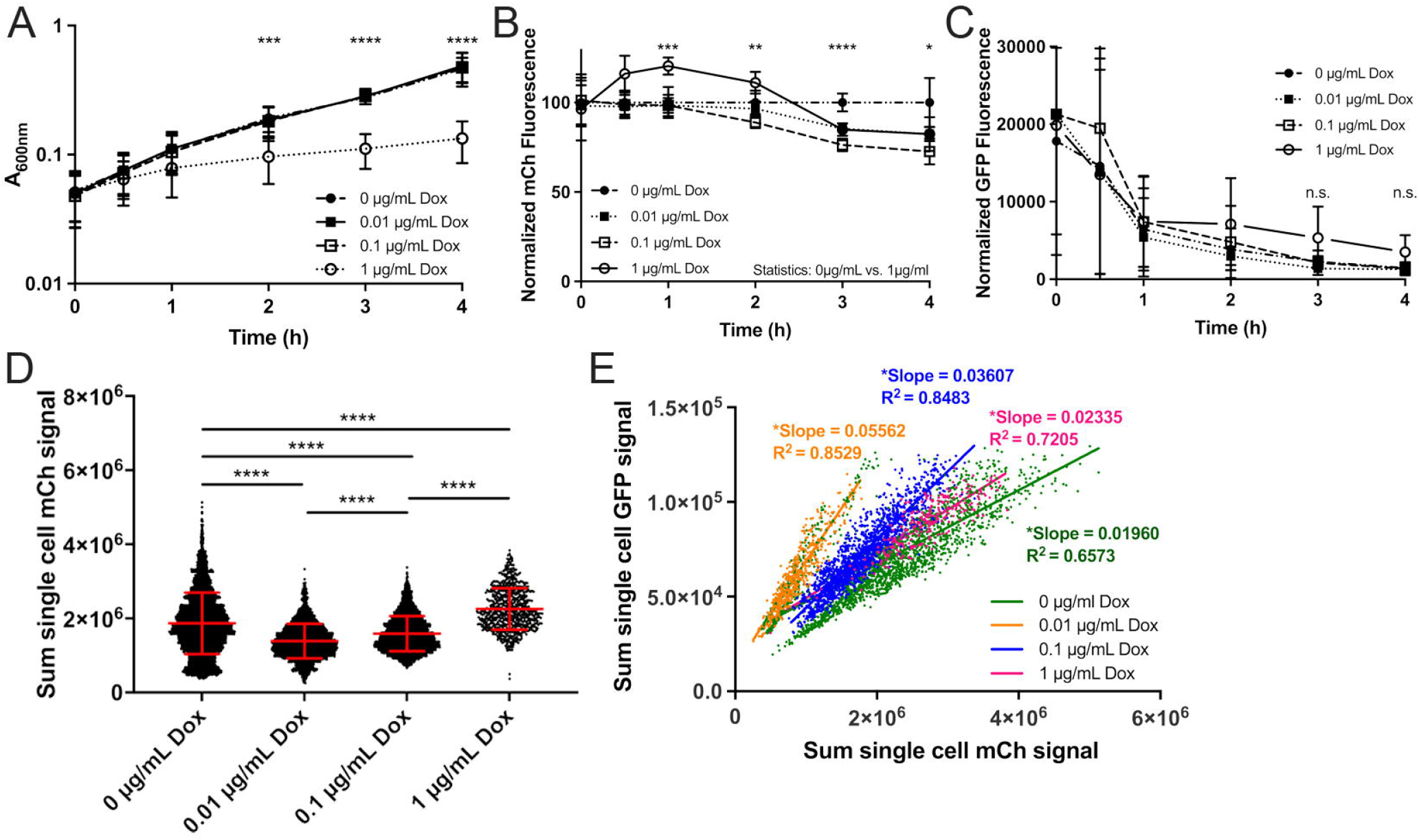
Growth inhibition with doxycycline promotes mCherry fluorescence accumulation. Exponential phase cultures of the revTetR-mCherry *dps* strain were treated with the indicated doses of doxycycline (Dox) at 0h (hours, h), aliquots were taken at the indicated timepoints to detect **(A)** optical density by absorbance (A_600nm_), **(B)** mCh fluorescence (normalized by dividing total fluorescence by OD values, untreated value set to 100%), and **(C)** GFP fluorescence normalized to OD. **(A-C)**: Data shown from 9 replicates. **(D)** Cells treated with the indicated doses of Dox for 4h were fixed for fluorescence microscopy. Sum single cell mCherry signal is shown. **(E)** Correlation plot depicting single cell sum mCherry and sum GFP signals. **(D-E):** Data shown from three biological replicates. Mean and standard deviation are shown. Statistics: **(A-C)** Two-way ANOVA, Tukey’s multiple comparison test, comparisons shown between untreated (0µg/ml) and 1µg/ml Dox; **(D)** Kruskal-Wallis one-way ANOVA with Dunn’s post-test; **(E)** linear regression with R^2^ value and slope of best-fit line, significantly non-zero slope indicates values are correlated. ****p<.0001, ***p<.001, **p<.01, *p<.05, n.s.: not-significant.

### Nitric oxide-stressed cells divide at a slower rate than unstressed cells

It has been shown recently that exposure to nitric oxide (NO) results in cell division arrest, through collapse of the FtsZ cell division machinery (48). Our results have also suggested NO-stressed *Yersinia pseudotuberculosis* preferentially survive doxycycline treatment during growth in the mouse spleen (22), which is consistent with a hypothesis that NO causes cell division arrest within host tissues, and subsequently promotes decreased antibiotic susceptibility. To test this hypothesis, we generated a strain containing the revTetR-mCherry reporter and *P_hmp::_gfp*, which can be used to mark NO-stressed cells (16, 17).

The revTetR-mCherry *P_hmp::_gfp* strain was treated with the slow-releasing NO donor compound DETA-NONOate for 2h, then further treated with ATc (1µg/ml). The *P_hmp::_gfp* reporter was used to detect NO exposure, and to mark the level of stress experienced by individual cells, which can be variable across the population in response to DETA-NONOate (16, 17). Growth inhibition during treatment with the NO donor, DETA-NONOate, was confirmed based on a significantly lower absorbance value in treated cultures at the 4h timepoint **(Figure 3A)**. Induction of *hmp* in response to the NO donor was also confirmed across all treatment groups **(Supplemental Figure 1)**. In the absence of ATc treatment, NO-exposed cells increased in mCherry signal over time, which is consistent with growth inhibition and decreased protein turnover **(Figure 3B)**. In cells treated with NO and ATc, the revTetR-mCherry fluorescence decreased between 0h and 2h, indicating some cell division within this timeframe, but signal slightly increased between 2h and 4h ATc treatment indicating cell division arrest **(Figure 3B)**. This timing was consistent with the growth inhibition observed at 4h in Figure 3A. In contrast, cells grown in the absence of NO treatment had a significant drop in mCherry fluorescence between 2h and 4h, suggesting significant levels of cell division events within this timeframe **(Figure 3B)**. Consistent with previous literature, these results indicate that NO treatment results in growth arrest, and reduced levels of cell division events (48), as detected by revTetR-mCherry.

**Figure 3.**
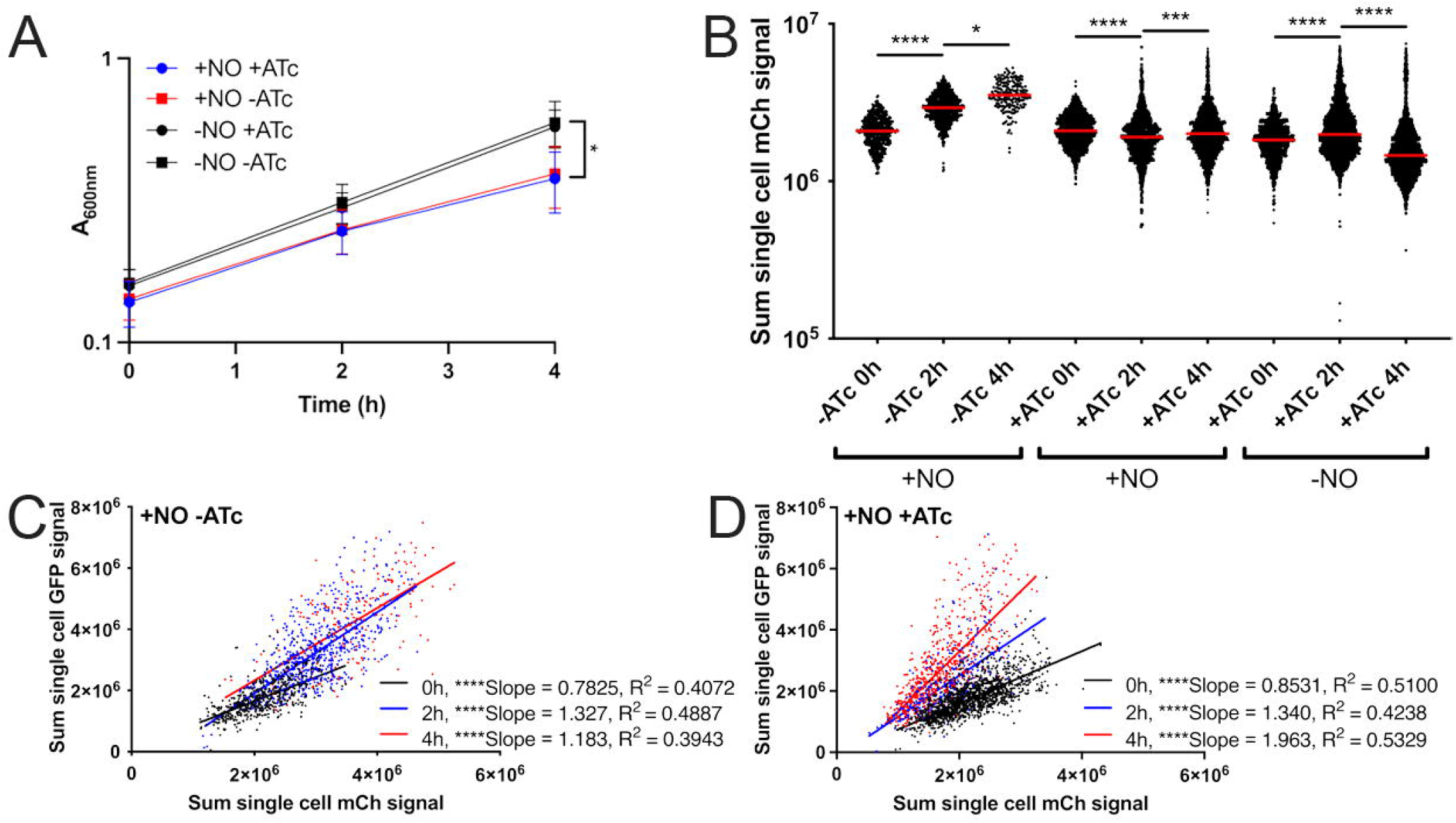
Nitric oxide-stressed cells divide at a slower rate than unstressed cells. Exponential phase cultures of the revTetR-mCherry *P_hmp_::gfp* strain were cultured two hours in the presence (+NO) or absence (-NO) of NO, then split, and either treated with 1µg/ml ATc (+ATc) or left untreated (-ATc). Timepoints are shown relative to ATc addition. **(A)** Aliquots were taken at the indicated timepoints to detect optical density by absorbance (A_600nm_). Mean and standard deviation are shown. **(B)** Aliquots of cells were fixed at the indicated timepoints for fluorescence microscopy. Sum single cell mCherry signal is shown. Median value/group is highlighted. **(C-E)** Correlation plots depicting single cell sum mCherry and mean GFP signals. Comparisons were made for NO-treated cells (+NO) in the presence (+ATc) and absence (-ATc) of ATc, cells were imaged at the indicated timepoints. Values were normalized to the max signal intensity detected in samples, represented by a value of 1.0. All data shown represents three biological replicates. Number of cells analyzed is shown. Statistics: **(A)** Two-way ANOVA, Tukey’s multiple comparison test, comparisons shown between untreated (-NO) and NO donor treated (+NO) samples; **(B)** Kruskal-Wallis one-way ANOVA with Dunn’s post-test; **(C-E)** linear regression with R^2^ value and slope of best-fit line, significantly non-zero slope indicates values are correlated. ****p<.0001, *p<.05, n.s.: not-significant.

To determine whether heightened NO stress correlated with a reduced level of cell division events, single cell mCherry (revTetR) and GFP (Hmp) values from NO-treated cells were compared in the absence and presence of ATc treatment using correlation plots and linear regression analyses. In the absence of ATc treatment, the distributions of cells within the NO-stressed bacterial populations were very similar in fluorescence across all timepoints **(Figure 3C)**. In the presence of ATc treatment, there was a clear shift in the NO-treated population towards reduced mCherry signal over time **(Figure 3D)**. This suggested some cell division still occurred in the presence of NO treatment, which was consistent with Figure 3B. However, the correlation between the two fluorescent signals strengthened over the timecourse based on increased R^2^ values **(Figure 3D)**. These data support the conclusion that cells experiencing higher level of NO stress are dividing more slowly, and retaining higher levels of revTetR-mCherry fluorescence.

### Low doses of doxycycline and anhydrotetracycline diffuse evenly across microcolonies and do not inhibit bacterial growth

One of the major goals for generating the revTetR reporter was to set-up a system to detect bacterial cell division events within host tissues over defined periods of time. To do this, we would need to administer tetracyclines during infection, ensure that the dosage used would not impact bacterial growth, and also diffuse evenly across microcolonies to allow for accurate detection of signal dilution and cell division. We have previously developed a model of doxycycline treatment of *Y. pseudotuberculosis* infection, and found that a single 40mg/kg injection of doxycycline significantly reduces bacterial CFUs and also accumulates at a higher concentration at the periphery of microcolonies (22). We hypothesized that lowering the tetracycline dose may allow us to administer sub-inhibitory levels of antibiotic that would still modulate the revTetR reporter, while allowing for more even diffusion of the antibiotic across microcolonies. Our rationale was that lower antibiotic concentrations would be more readily eliminated from circulation, and so antibiotic movement into the spleen would occur within a more defined timeframe, limiting continued accumulation.

To identify a tetracycline concentration that results in equal diffusion across microcolonies, a previously characterized reporter, *P_tetA_::mCherry-ssrA* (TetON), was used for tetracycline detection (22). The TetON reporter contains the wild-type *tetR* gene; addition of tetracyclines induce the production of mCherry, and therefore levels of mCherry can be used as a readout for antibiotic exposure **(Figure 4A)**. C57BL/6 mice were inoculated intravenously with the TetON *gfp*^+^ strain, which has constitutive expression of GFP alongside the TetON reporter, allowing for visualization of relative reporter expression. A single dose of 4mg/kg Dox or 4mg/kg ATc was administered intraperitoneally at 48 hours post-inoculation. This timepoint was chosen for antibiotic administration since we can detect subsets of bacteria responding to host-derived nitric oxide at this timepoint, thus allowing us to ask whether stressed subpopulations of bacteria are dividing more slowly than unstressed bacteria.

**Figure 4.**
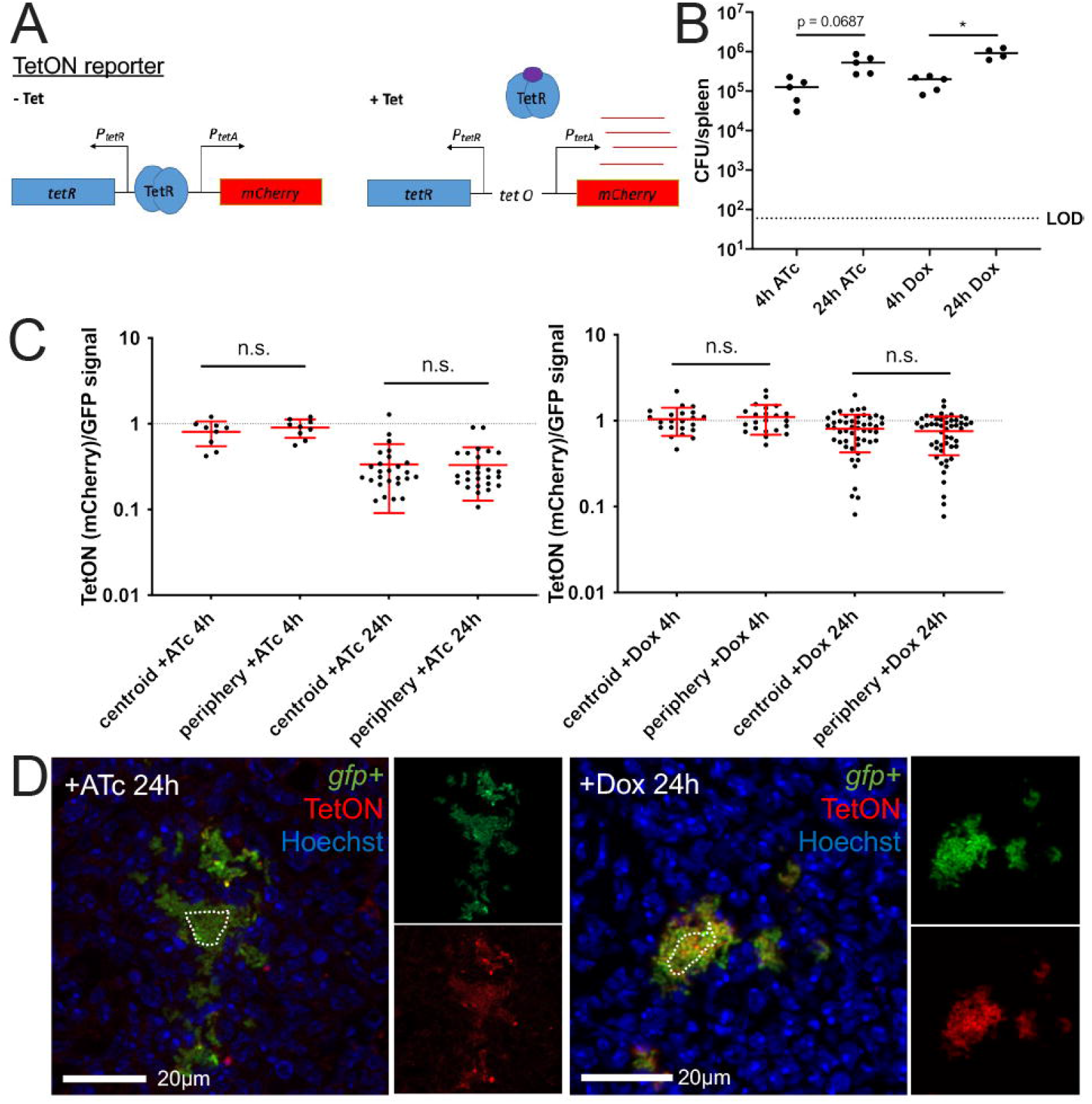
Low doses of doxycycline and anhydrotetracycline diffuse evenly across microcolonies and do not inhibit bacterial growth. C57BL/6 mice were infected intravenously with the TetON *gfp^+^* strain and a single dose of Dox (4mg/kg) or ATc (4 mg/kg) was administered intraperitoneally at 48h post-inoculation. Spleens were harvested at the indicated timepoints after treatment. **(A)** Schematic of the TetON reporter: addition of tetracycline relieves TetR repression and allows for *mCherry* transcription. **(B)** CFU/spleen quantification at the indicated timepoints post-treatment. Dotted line: limit of detection, dots: individual mice. **(C)** Reporter signal was detected with fluorescence microscopy and is shown as the ratio of TetON (mCherry)/GFP signal at the centroid and periphery of microcolonies. Timepoints indicate time post-treatment with ATc (left panel) or Dox (right panel). Dotted line: equivalent levels of mCherry and GFP signal. Each dot: individual microcolony. Mean and standard deviation are shown. **(D)** Representative images. Merged images and individual channels are shown. Cells outside the dotted line are defined as peripheral. Dataset represents 4-5 mice/group, collected from two independent experiments; 2-8 microcolonies were analyzed/spleen from a single cross-section of tissue. Statistics: **(B)** Kruskal-Wallis one-way ANOVA with Dunn’s post-test; **(C)** Wilcoxon matched-pairs; *p<.05, n.s.: not-significant.

At 4h or 24h post-treatment, spleens were harvested to quantify CFUs and quantify reporter expression by fluorescence microscopy. Bacterial numbers continued to increase between 4h and 24h post-treatment with 4mg/kg doses of either ATc or Dox, confirming this dosage does not interfere with bacterial growth **(Figure 4B)**. Microcolonies were then visualized by fluorescence microscopy to determine if ATc and Dox diffuse evenly across microcolonies at this dosage. The centroid and periphery of each microcolony were quantified for mCherry and GFP signal, and the ratio of mCherry/GFP was graphed. There were no significant differences in mCherry/GFP ratios between the centroid and periphery of microcolonies treated with either ATc or Dox, suggesting that each drug was diffusing evenly across microcolonies **(Figure 4C)**. The mCherry/GFP ratios remained comparable at the centroid and peripheries even at 24 hours after treatment, showing that throughout the infection time course, there was even diffusion of the drugs **(Figure 4C, 4D)**. These data here suggest that lowering the tetracycline dosage 10-fold results in even diffusion of a sub-inhibitory dose of tetracyclines, which will allow us to modulate revTetR reporter expression in all cells across the microcolony without impacting bacterial growth.

### revTetR signal appears to increase in both the centroid and periphery of microcolonies

To determine if there are slowly-dividing subpopulations of bacteria within microcolonies, we infected mice with the revTetR-mCherry *gfp*^+^ strain, which constitutively expressed *gfp*, and treated mice with ATc at 48h post-inoculation or left mice untreated. Mice were sacrificed at either 4h or 24h post-treatment, and spleens were harvested to quantify CFUs and visualize reporter expression by fluorescence microscopy. Bacterial CFUs were comparable between ATc-treated and untreated mice at both timepoints, suggesting that ATc treatment did not impair bacterial growth **(Figure 5A)**. However, we did not see an increase in CFUs between 4h and 24h, which suggests there was limited, or variable, bacterial growth within this selected timeframe for this bacterial strain. To determine if there were differences in reporter expression, mCherry signal was quantified relative to the constitutive GFP signal at the periphery and centroid of individual microcolonies. Since we have previously shown that bacteria experience heightened levels of NO stress and express higher levels of the T3SS at the periphery of microcolonies, we were expecting to see heightened levels of mCherry signal relative to GFP at the periphery of microcolonies. At 4h post-treatment, we did not observe any differences in mCherry signal at the periphery of microcolonies compared to the centroid, in the absence or presence of ATc treatment **(Figure 5B)**. This suggested there may not have been many cell division events within the first 4h of treatment, which is why tissue was also harvested 24h post-treatment. At 24h post-treatment, untreated mice had significantly lower levels of mCherry signal at the periphery of microcolonies, and ATc treated mice also had significantly lower mCherry signal at the periphery, indicating we were not seeing increased mCherry signal at the periphery of microcolonies as expected **(Figure 5C, 5D)**. It did appear that ATc treated microcolonies had higher mCherry signals at both the centroid and periphery relative to untreated microcolonies, although this was not statistically significant. These results suggest that revTetR signal remained high after ATc treatment, indicating there may be slowly dividing cells present within microcolonies. Without much bacterial growth across these timepoints in general, it was difficult to assess whether slow-growing cells were present in a specific spatial location or part of a subpopulation without an additional reporter alongside revTetR.

**Figure 5.**
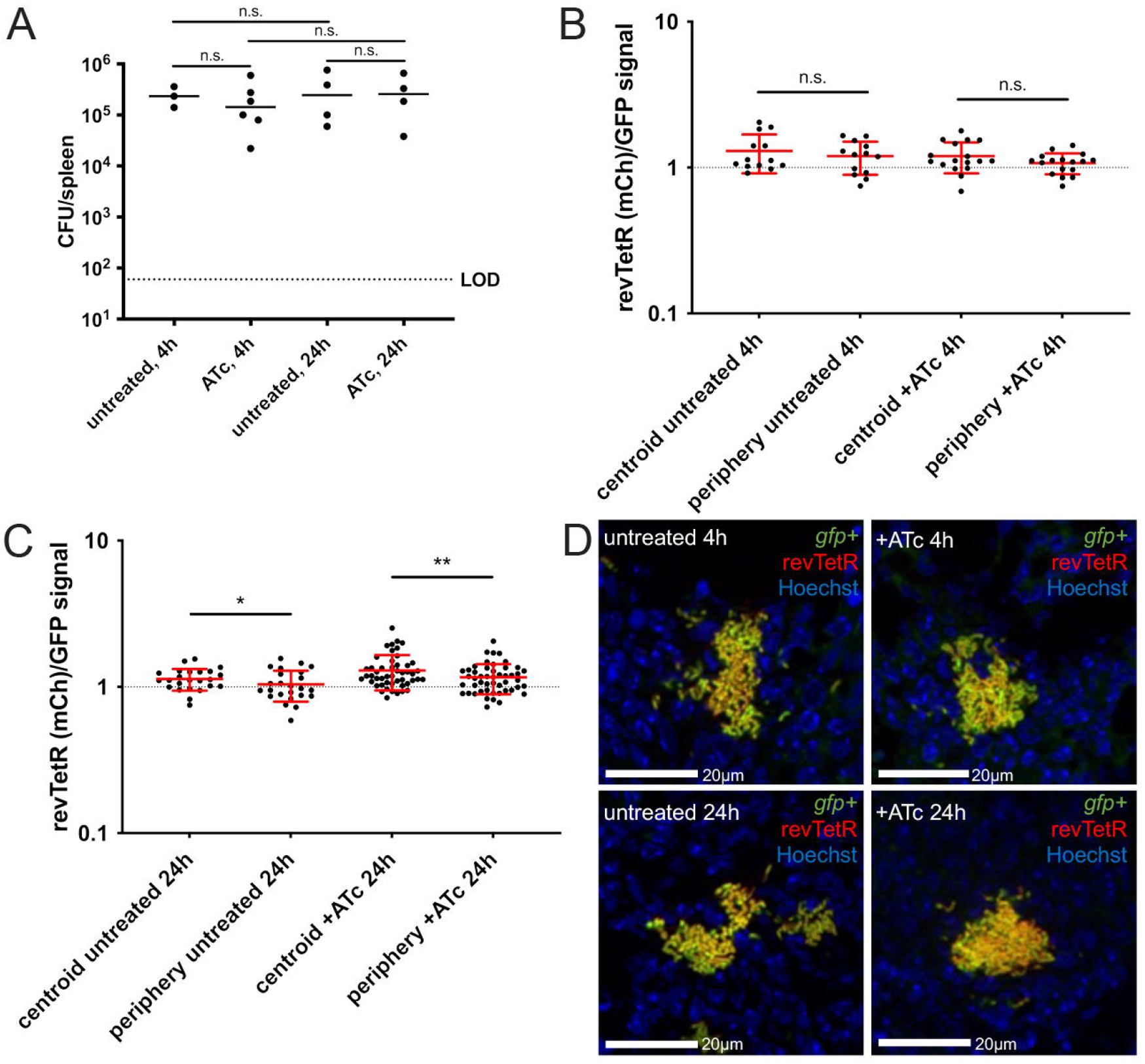
revTetR-mCherry signal appears to increase in both the centroid and periphery of microcolonies. C57BL/6 mice were infected intravenously with the revTetR-mCherry *gfp^+^* strain and a single dose of ATc (4 mg/kg) was administered intraperitoneally at 48h post-inoculation. Spleens were harvested at the indicated timepoints after treatment to quantify CFUs and prepare tissue for fluorescence microscopy. **(A)** CFU/spleen quantification at the indicated timepoints post-treatment. Dotted line: limit of detection, dots: individual mice, horizontal lines: median values. **(B)** Reporter signal at 4h quantified using the ratio of revTetR (mCherry)/GFP signal at the centroid and periphery of microcolonies. Dotted line: equivalent levels of mCherry and GFP signal. Each dot: individual microcolony. Mean and standard deviation are shown. **(C)** Reporter signal at 24h quantified using ratio of revTetR (mCherry)/GFP signal at the centroid and periphery of microcolonies. Mean and standard deviation are shown. **(D)** Representative images. Dataset represents 4-6 mice/group, collected from three independent experiments; 2-13 microcolonies were analyzed/spleen from a single cross-section of tissue. Statistics: **(A)** Kruskal-Wallis one-way ANOVA with Dunn’s post-test; **(B-C)** Wilcoxon matched-pairs; **p<.01, *p<.05, n.s.: not-significant.

### NO stress correlates with heightened mCherry signal accumulation and slowed cell division rates

Hmp^+^ bacteria responding to NO stress preferentially survive doxycycline treatment, suggesting they may represent a slow-growing subpopulation (22). Consistent with this, in culture, NO-stressed cells accumulated heightened mCherry signal, suggesting they were dividing more slowly than unstressed cells **(Figure 3)**. However, in our mouse model it was difficult to determine if peripheral cells of the microcolony had heightened revTetR signal without marking the peripheral subpopulation with the *hmp* reporter. To determine if NO-stressed bacteria are dividing more slowly than unstressed cells within host tissues, we infected mice with the revTetR-mCherry *P_hmp_::gfp* strain and treated mice with ATc at 48h post-inoculation, or left mice untreated. Spleens were harvested at 4h and 24h post-treatment to quantify CFUs and visualize reporter expression by fluorescence microscopy. CFUs within ATc-treated mice did not differ significantly from CFUs within untreated mice; again, we noted that there was not significant growth based on total CFUs between 4h and 24h, but anticipated that there would be sufficient replication at the level of individual microcolonies to assess differences in revTetR signal **(Figure 6A)**.

**Figure 6.**
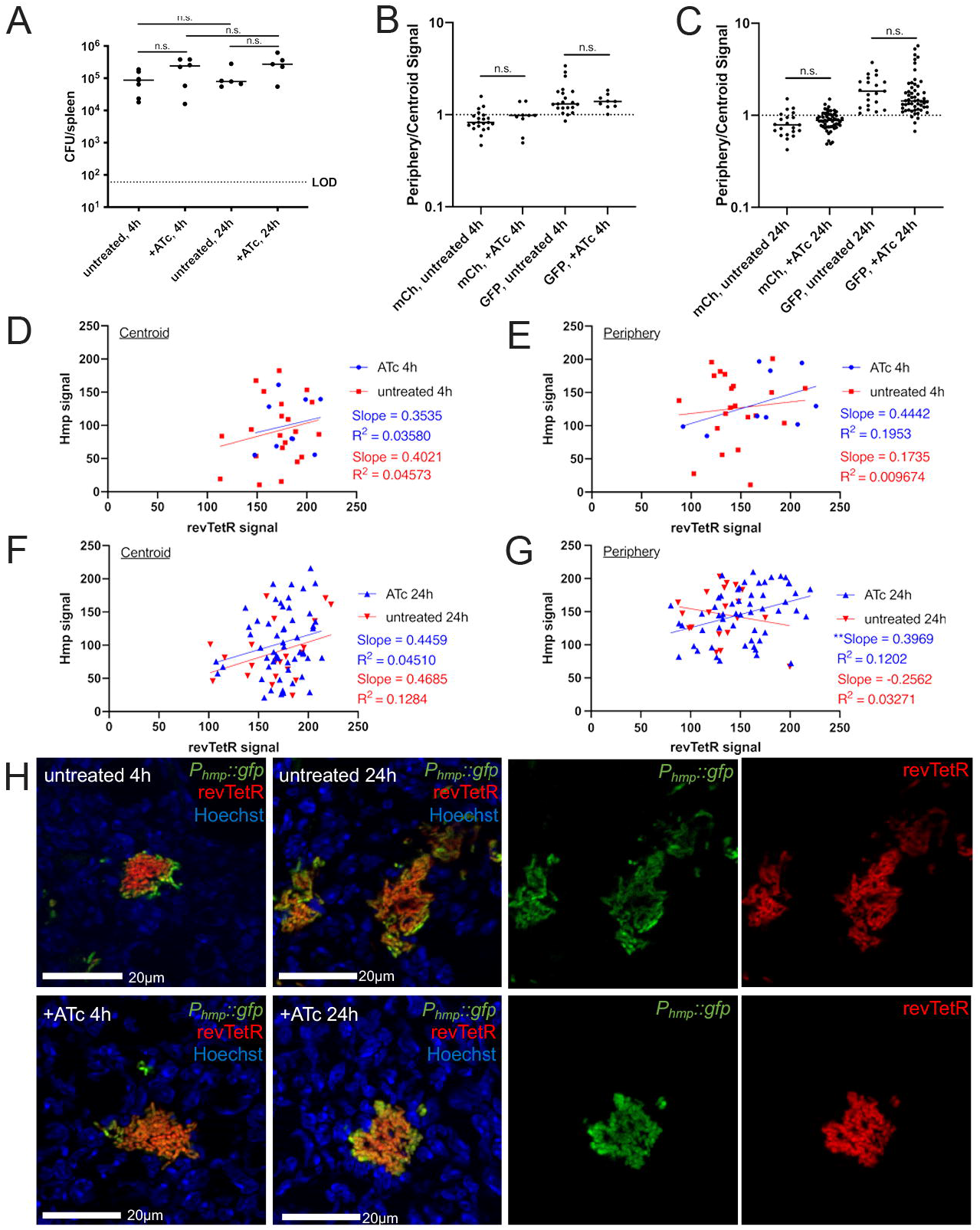
NO stress correlates with heightened mCherry signal accumulation and slowed cell division rates. Mice were infected with the revTetR-mCherry *P_hmp_::gfp* strain and a single dose of ATc (4 mg/kg) was administered intraperitoneally at 48h post-inoculation. Spleens were harvested at the indicated timepoints after treatment and prepared for fluorescence microscopy. **(A)** CFU/spleen quantification. Dotted line: limit of detection, dots: individual mice. **(B)** Reporter signal at 4h quantified using ratio of periphery/centroid signal for either mCherry (mCh) or GFP. Dotted line: equivalent levels of mCherry and GFP signal. Each dot: individual microcolony. **(C)** Reporter signal at 24h quantified using ratio of periphery/centroid signal for either mCh or GFP. **(D-G)** Correlation plots depicting single cell revTetR signal (mCherry) and Hmp signal (GFP) at the indicated timepoints, comparing +/- ATc treatment. Graphs represent either centroid or peripheral values, dots: individual microcolonies. **(H)** Representative images with the green and red channels shown alongside merged images for 24h post-treatment images. Dataset represents 5-6 mice/group, collected from three independent experiments; 3-8 microcolonies were analyzed/spleen from a single cross-section of tissue. Median values shown with horizontal lines. Statistics: (A-C) Kruskal-Wallis one-way ANOVA with Dunn’s post-test **(D-G)** linear regression with R^2^ value and slope of best-fit line, significantly non-zero slope indicates values are correlated. **p<.01, n.s.: not-significant.

Periphery and centroid measurements were taken from each microcolony image and combined into a single periphery/centroid ratio for each microcolony to allow for comparison across treatment groups. At 4h post-ATc treatment, the average ratio for the *hmp* reporter signal was above a value of 1, indicating higher expression in the peripheral cells **(Figure 6B)**. The average ratio for revTetR-mCherry values was close to 1 in both ATc-treated and untreated mice, suggesting equal expression in peripheral and centroid cells and very little cell division in this time frame **(Figure 6B)**. At 24h post-ATc treatment, GFP ratios remained above 1, indicating high peripheral expression of the *hmp* reporter **(Figure 6C)**. The revTetR-mCherry ratio appeared to increase slightly, but remained below a value of 1, and was not significantly different from untreated mice **(Figure 6C)**. This again suggested there was little cell division within this timeframe, since we could not detect much impact of the ATc treatment. We then used linear regressions to determine if there was a correlation in fluorescent signals at distinct spatial locations within the microcolony, and overall saw little impact of the ATc treatment on the fluorescence of the cells **(Figure 6D-H)**. The exception was the analysis at the periphery 24h post-treatment, where ATc treatment resulted in a slight positive correlation suggesting Hmp^+^ cells were less likely to divide **(Figure 6G, 6H)**. The limited levels of cell division with this strain *in vivo* made these results difficult to interpret, and we then sought to modify the reporter to generate a new revTetR construct with normal growth kinetics *in vivo*.

### Dilution of the revTetR-YFP signal coincides with the replication of fast-growing cells

To address the growth issues in the revTetR-mCherry strain evident in previous experiments, we modified the revTetR reporter to express YFP instead of mCherry (*revtetR::P_tetA_::yfp)*. We surmised that the growth issues were due to the more toxic and disruptive nature of constant mCherry overexpression in the absence of tetracycline derivatives, and YFP is known to be less toxic for cells (49, 50).

To determine if revTetR-YFP strains exhibit normal growth kinetics, and confirm NO stress is sufficient to slow growth and promote YFP signal accumulation, we generated a strain containing the revTetR-YFP reporter and *P_hmp_::mCherry* (*revtetR::P_tetA_::yfp P_hmp_::mCherry*). Cells were grown in the presence and absence of the NO donor compound DETA-NONOate for 2h, then treated with ATc (1µg/ml) as described above. Aliquots were taken at the indicated timepoints after ATc addition to quantify single cell fluorescence by microscopy. Growth inhibition due to the NO donor was confirmed by negligible signal dilution between 0h and 2h post-ATc treatment **(Supplemental Figure 2A)**, and all treated cultures responded to the NO donor based on induction of *hmp* **(Supplemental Figure 2B)**. Cells exposed to NO and ATc had a slight signal dilution between 0-2h and 2-4h, suggesting some continued cell division, however cells grown in the absence of NO showed dramatic YFP signal dilution between 0-2h **(Supplemental Figure 2A)**. This suggested that the revTetR-YFP construct behaves similarly to revTetR-mCherry, and can be used to mark growth-arrested cells. However, unstressed cells (- NO, +ATc) no longer had signal dilution between 2-4h post-ATc treatment, suggesting growth was slowing in these cultures. Since cells were incubated with NO for 2h before ATc treatment, this indicates that the cells were beginning to enter stationary phase by the 4h timepoint (6h of growth). These changes in growth kinetics indicate that the revTetR-YFP strain may grow faster than the revTetR-mCherry strain in culture. Additionally, the signal dilution between 2-4h in the +NO +ATc group **(Supplemental Figure 2A)** suggests cells are recovering from NO exposure and are starting to replicate after 6h exposure. This is consistent with our previous work (16), and consistent with the presence of some Hmp^low^/mCherry^low^ cells in the +NO +ATc group at 4h **(Supplemental Figure 2B)**.

To ensure we were capturing the timeframe of growth arrest, we modified the *in vitro* NO donor experiment to shorten the NO exposure. Cells containing the revTetR-YFP reporter and *P_hmp_::mCherry* were treated with the NO donor compound for only 30 minutes before ATc addition, and single cell fluorescence was quantified via microscopy. Cells grown in the absence of ATc treatment continued to accumulate YFP signal, which was unsurprising **(Supplemental Figure 3A)**. Heightened *hmp* induction in the +NO groups compared to the -NO groups confirmed NO exposure throughout the timecourse **(Supplemental Figure 3B)**. With this experimental protocol, cells exposed to the NO donor and ATc treatment increased in revTetR-YFP signal between 0-2h, and only a slight decline between 2-4h, indicating overall growth arrest **(Figure 7A)**. Cells grown in the absence of the NO donor with ATc treatment showed continuous signal dilution, indicating continuous growth and replication **(Figure 7A)**. These results suggest that in this faster growing strain, YFP signal dilution does correspond with bacterial cell growth, and signal accumulation corresponds with growth arrest.

**Figure 7.**
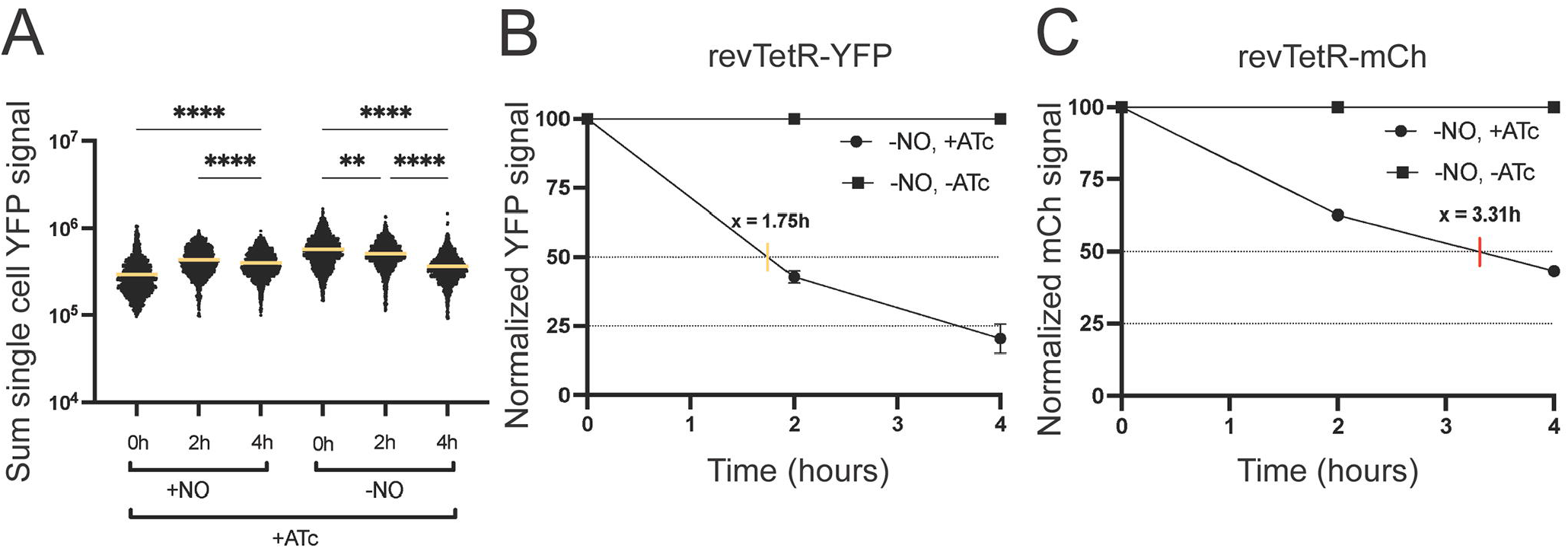
Dilution of the revTetR-YFP signal coincides with replication of fast-growing cells. The revTetR-YFP *P_hmp_::mCherry* strain was grown in the presence (+NO) or absence (- NO) of NO for 30 min. Cultures were then treated with 1µg/ml ATc (+ATc) or left untreated. Aliquots were taken at timepoints after ATc addition. **(A)** Sum single cell YFP signal determined by microscopy. Bars: mean. At least 1,500 individual cells were quantified per group. **(B-C)** Replication rates of unstressed (-NO) cells were calculated by first normalizing to fluorescent signal at 0h, then values of treated cells (+ATc) were further normalized to the untreated values (-ATc, set to 100%) to depict signal dilution over time. Linear regressions were used to calculate the time it takes for 100% of the cells in culture to undergo one full replication (50% dilution of the signal). This was done for **(B)** revTetR-YFP (yellow tick, 1.75h for full replication cycle) and **(C)** revTetR-mCherry (red tick, 3.31h). Statistics: **(A)** Kruskal-Wallis one-way ANOVA with Dunn’s post-test; ****p<.0001, **p<.001.

Since our results above indicate the growth kinetics are different between the revTetR-YFP and revTetR-mCherry strains, we estimated the length of time it takes for 100% of the cells in culture to undergo one full replication cycle, as indicated by a 50% dilution of the revTetR signals. Calculations were completed using data from unstressed cells: -NO +ATc and -NO -ATc groups. YFP signals from our single cell microscopy data were normalized to 100% at 0h, then the ATc-treated group (-NO +ATc) data was normalized to the untreated control group (-NO - ATc) to account for signal dilution over time. Using a linear regression equation, the revTetR-YFP strain was estimated to undergo one replication cycle, or doubling, in 1.75h **(Figure 7B)**. This is consistent with normal growth kinetics of *Y. pseudotuberculosis* at 37° C (16). We then confirmed that the revTetR-mCherry strain does indeed grow slower, with an estimated 3.31h for one replication cycle **(Figure 7C)**. Together, these data indicate that the revTetR-YFP reporter can be used to differentiate between dividing and non-dividing cells, and confirm the growth issues seen in the revTetR-mCherry strains have been resolved with revTetR-YFP.

### Cells that lack NO stress rapidly dilute revTetR-YFP signal indicating faster cell division rates

Our previous work indicates that cells at the periphery of the microcolony respond to NO stress and may be slow-growing, since Hmp^+^ cells had reduced susceptibility to a ribosome-targeting antibiotic (22). While our *in vitro* data support this **(Figure 7)**, our previous study using a modified TIMER protein (DsRed_42_) to detect slow-growing bacteria suggested that NO stress is transient within host tissues (16). At late timepoints during infection, Hmp^-^ cells at the centroid of microcolonies were instead slow-growing due to nutrient limitation (16). In our mouse infections with the revTetR-mCherry *P_hmp_::gfp* strain above, it appeared that cells responding to NO stress (Hmp^+^, GFP^+^) may have higher mCherry levels **(Figure 6G)**. Due to growth issues with the strain, it was difficult to interpret these results. To clarify if the subset of NO-exposed cells are truly growing at a slower rate than unstressed cells within host tissues, we infected mice with the revTetR-YFP *P_hmp_::mCherry* strain, then treated with ATc (+ATc) at 48h post-inoculation, or left mice untreated (unt). We confirmed the revTetR-YFP *P_hmp_::mCherry* strain was growing well within host tissues based on significant increases in CFUs in the spleen, comparing between 4h and 24h post-ATc treatment **(Figure 8A)**. Spleens were processed for fluorescence microscopy, periphery and centroid measurements were taken from each microcolony, and combined into a single periphery/centroid ratio to allow for comparison across treatment groups. At 4h post-treatment, the average ratio for the *hmp* (mCh) reporter signal was above a value of 1 in both untreated (unt) and treated mice (+ATc), indicating higher expression in the peripheral cells, as expected. The average ratio of revTetR-YFP signal was close to 1 in untreated mice, suggesting equal expression in peripheral and centroid cells **(Figure 8B)**. In contrast, ATc-treated mice had a ratio above 1, suggesting there is higher YFP expression in the periphery relative to the centroid **(Figure 8B)**. Similar trends held true at 24h post-treatment, although the *hmp* (mCh) reporter signal was higher in treated mice for unknown reasons **(Figure 8C)**.

**Figure 8.**
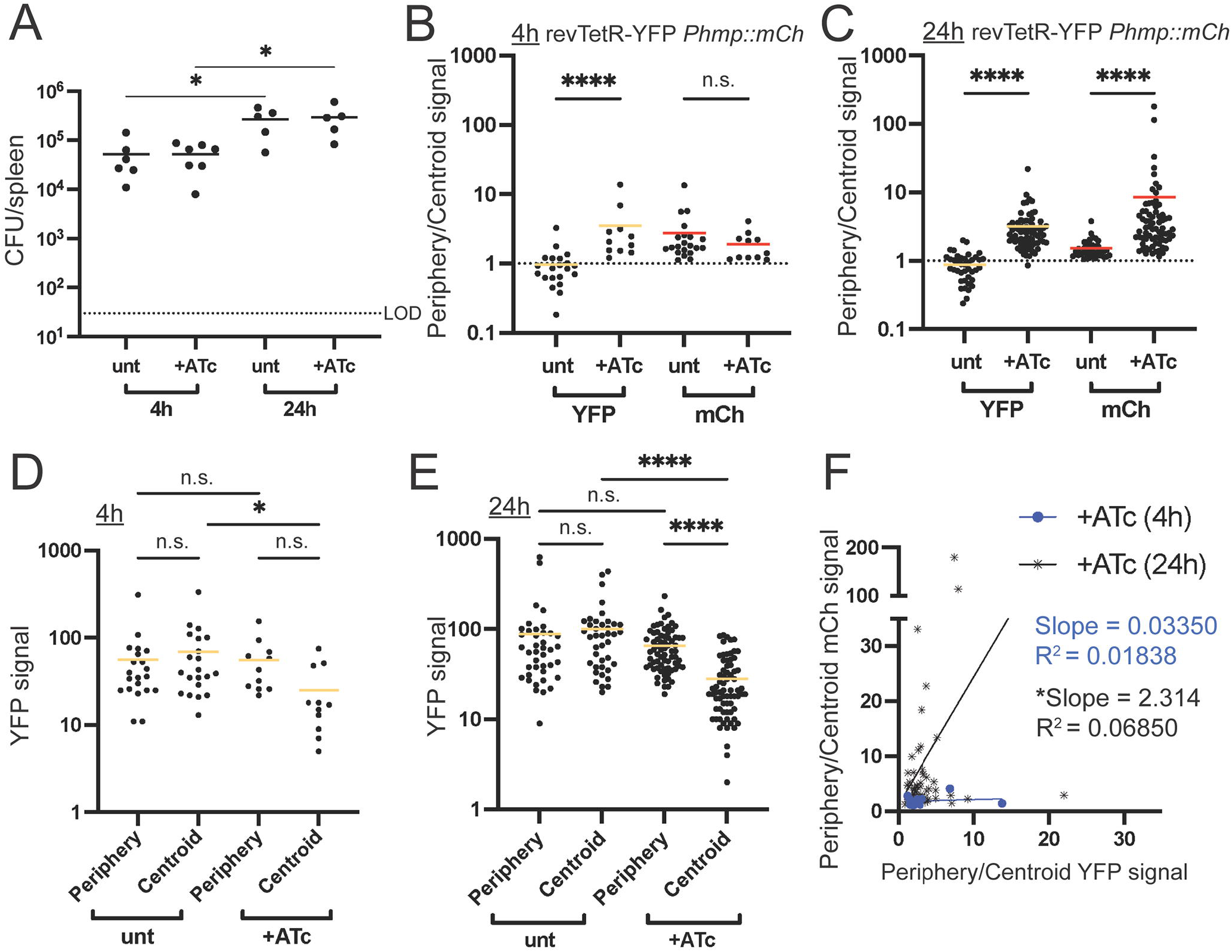
Cells that lack NO stress rapidly dilute revTetR-YFP signal indicating faster cell division rates. Mice were infected with the revTetR-YFP *P_hmp_::mCherry* strain, then either injected with 4 mg/kg ATc (+ATc) or left untreated (unt) at 48h post-inoculation. **(A)** CFU/spleen was quantified for each mouse at the indicated timepoints post-ATc treatment. Dotted line indicates limit of detection (LOD), and each dot represents one mouse. **(B-C)** The ratio of periphery/centroid signals were quantified with revTetR (YFP) and Hmp reporter (mCherry) signals at **(B)** 4h and **(C)** 24h post-ATc treatment, along with the raw YFP signals at **(D)** 4h and **(E)** 24h post-treatment. **(F)** Correlation plots are shown for periphery/centroid ratios for revTetR (YFP) and Hmp (mCherry) reporter signals, comparing treated groups at each timepoint. **(B-F)** Each dot represents a microcolony. Mean values are highlighted. Data represent 4-5 mice per group per timepoint, with 11-71 microcolonies analyzed for each group and timepoint. Statistics: **(A-E)** Kruskal-Wallis one-way ANOVA with Dunn’s post-test; **(F)** linear regression was used to determine correlation between values, a significant deviation from zero indicates the values are correlated; ****p<.0001, *p<.05.

An increased periphery/centroid ratio can mean either the periphery signal increased, or conversely, the centroid signal decreased. Because of the nature of fluorescence dilution, we would expect this means the centroid is diluting revTetR-YFP signal more rapidly than the periphery, suggesting the centroid is more rapidly dividing. To assess this, raw YFP signal values were compared across treatment groups and timepoints. At 4h post-ATc treatment, YFP signals for untreated mice were not significantly different between the centroid and periphery **(Figure 8D)**. The YFP signal in the centroid of ATc-treated mice was trending, but not significantly lower than the periphery **(Figure 8D)**. However, the centroids of ATc-treated mice were significantly lower than the centroids of untreated mice, suggesting cell division and signal dilution at the centroid **(Figure 8D)**. At 24h post-ATc treatment, untreated mice had similar levels of YFP signal at the centroid and periphery of microcolonies, while centroids of treated mice had significantly lower levels of YFP when compared to treated peripheries or untreated centroids **(Figure 8E)**. These results suggest significant cell division within the centroid of microcolonies, and relative growth arrest at the periphery, since peripheral cells were not changing in YFP signal **(Figure 8E)**. These conclusions are corroborated by the correlation of revTetR-YFP and mCherry (Hmp) periphery/centroid ratios, where revTetR and Hmp exhibit a weak correlation at 24h post-treatment, based on a significantly non-zero slope **(Figure 8F)**. It is important to note that while cells in the periphery tend to have higher YFP signals in the treated mice at 24h **(Figure 8F)**, some subsets of peripheral cells do not exhibit this high YFP signal **(Figure 9)**. It is, however, striking, how much YFP dilution occurs at the centroid of microcolonies, which strongly supports the conclusion that the centroid is rapidly dividing **(Figure 8E, 9)**. Overall, this microcolony phenotype is consistent with our previous findings that only small subsets of peripheral cells actively respond to NO stress at any given timepoint (16), which appears to correlate with detection of growth arrest with revTetR-YFP.

**Figure 9.**
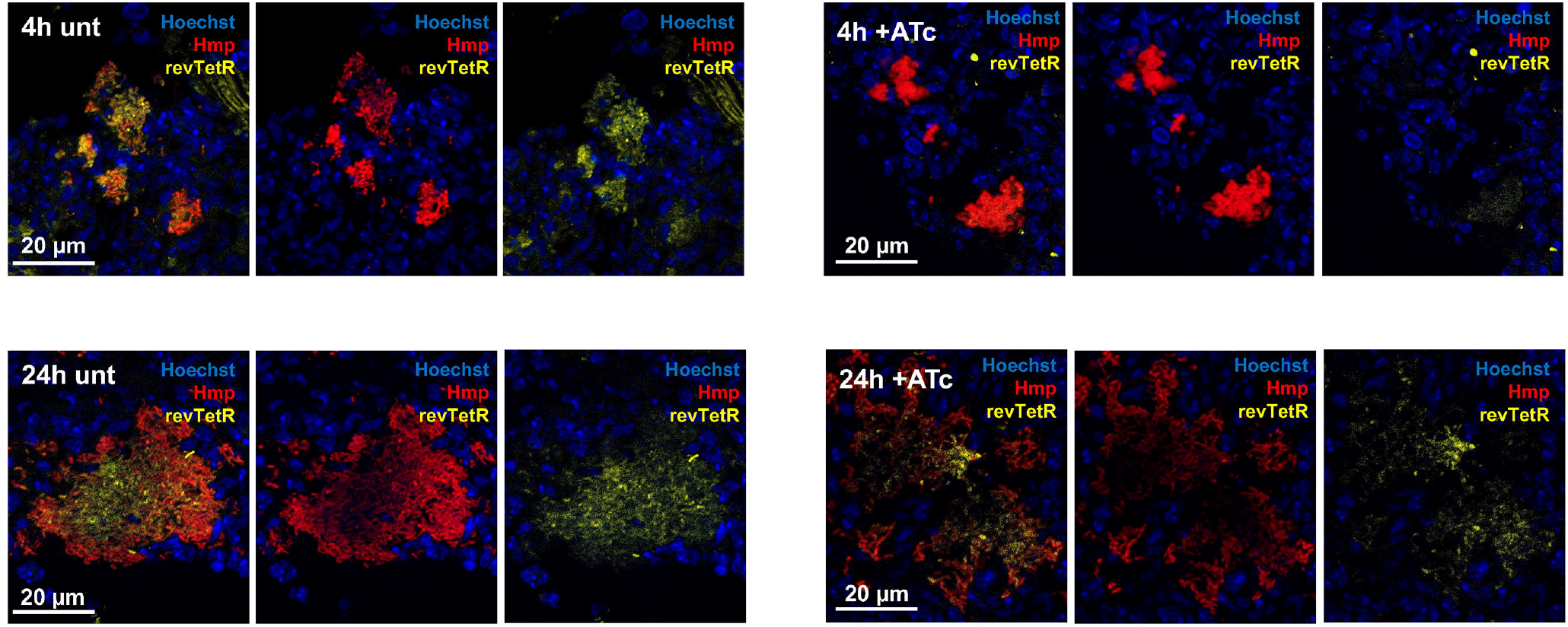
Representative images of spleens during revTetR-YFP *P_hmp_::mCherry* infection. Representative images are shown from spleens of mice infected with the revTetR-YFP *P_hmp_::mCherry* strain, as described and quantified in Figure 8. Merged images are shown alongside red and yellow single channels images at the indicated timepoints after ATc treatment.

## Discussion

In this study, we have developed revTetR constructs that can be utilized to detect bacterial cell division events within host tissues using a fluorescence dilution approach. We have shown both revTetR-mCherry and revTetR-YFP constructs result in heightened fluorescent signal in growth-arrested cells in culture, and have also shown that nitric oxide (NO) stress is sufficient to promote increased fluorescent signal in culture **(Figures 3, 7)**. This was consistent with previous studies showing NO exposure results in an arrest of cell division (48). Our *in vivo* results in the mouse model of infection were initially less clear; our revTetR-mCherry strains had notable slowed growth within host tissues **(Figures 5A, 6A)**, which made it difficult to differentiate dividing and non-dividing cells. Experiments with revTetR-mCherry alongside a *hmp* reporter hinted that Hmp^+^ cells may be slowly dividing **(Figure 6G)**, but this conclusion remained unclear until experiments were completed with the revTetR-YFP *P_hmp_::mCherry* strain. Using revTetR-YFP, we detected clear differences in cell division rates across microcolonies. Bacterial cells at the centroid are clearly replicating more quickly and diluting YFP signal, while we saw YFP signal was maintained within peripheral cells **(Figures 7D, 7E)**. Some Hmp^+^ cells retained YFP signal while others did not, which we believe reflects transient NO exposures within host tissues (16).

The Hmp^+^ subpopulation was a focus of this study primarily because we knew this subpopulation preferentially survives doxycycline treatment in the *Y. pseudotuberculosis* mouse model of infection. We hypothesized this was due to increased levels of growth arrest as a result of NO exposure (22, 48). However, it is important to note that the Hmp^+^ subpopulation is a complex mixture of cells, that also includes subsets expressing high levels of the type-III secretion system (T3SS) due to direct neutrophil contact (17). Based on the pattern of revTetR-YFP signal within host tissues **(Figure 9)**, revTetR-YFP more closely correlates with Hmp reporter expression than T3SS, but additional experiments could directly assess this. We also chose to mark the Hmp^+^ subpopulation here with a stable fluorescent protein to detect NO exposure. However, we know that only subsets of this population are actively responding to NO at any given timepoint (16), which adds further complexity to this subpopulation. It will be very interesting to utilize this revTetR-YFP construct in future experiments to isolate bacterial cells with slowed division rates from the host environment, and further characterize the pathways that are associated with bacterial slowed growth within host tissues.

Importantly, we believe the cell division rates within microcolonies vary over the course of infection, and revTetR-YFP will allow us to probe this in more detail by selecting different timepoints over the course of infection for ATc treatment and infection endpoints. Here, we chose to compare between 52h and 72h post-inoculation, which equates to +ATc 4h and 24h, respectively. These were timepoints where we expected rapid expansion within the bacterial population, allowing us to differentiate between slowly-dividing and quickly-dividing cells. These timepoints worked well for distinguishing differential growth rates with revTetR-YFP, and showed the centroid of microcolonies were dividing rapidly and diluting fluorescent signal **(Figure 9)**. However, this does seem to contrast with some of our other results that indicate the centroid is slow-growing (16). In Patel P, *et al.*, slowed growth was detected in the centroid of microcolonies with a slow-folding DsRed variant, DsRed_42_, specifically at 72h post-inoculation (16). We hypothesize the timepoint is key here, and with the timepoints we have chosen in this study, we are capturing a period of rapid growth that precedes nutrient depletion at later timepoints. By switching from revTetR-mCherry to revTetR-YFP, we have generated strains without growth attenuation, and also generated a construct where revTetR-YFP fluorescence could be directly compared with DsRed_42_ to better understand bacterial growth dynamics over the course of infection.

Experiments with the revTetR-mCherry constructs have shown us that to truly detect differences in cell division rates *in vivo*, we need to sample timepoints where there is rapid expansion within the bacterial population. Based on our estimates in Figure 7, revTetR-YFP strains double approximately every 1.75h during culture at 37° C, while revTetR-mCherry strains double approximately every 3.3h, and may take as long as 4h to divide, based on Figure 1D. The doubling time, or replication rate, thus essentially establishes the sensitivity of the system, and needs to be taken into account when choosing the timepoints for experiments. Conversely, by completing full timecourses, researchers would be able to utilize these reporters to approximate the growth rate of strains grown under different conditions. We believe this application could be particularly useful in approximating the bacterial cell division rates within host tissues, and plan to explore this in future experiments.

In this study, we compared revTetR-mCherry reporter signal during treatment with two tetracycline derivatives, the active antibiotic, doxycycline (Dox), and the inactive derivative, anhydrotetracycline (ATc). Both tetracycline derivatives bind to TetR and revTetR, however ATc is predicted to bind these proteins with higher affinity, and has poor affinity to bacterial ribosomes (46, 47). This allowed us to modulate revTetR with ATc without significantly inhibiting translation, however we would expect to see translational inhibition during Dox treatment, and did see evidence of this when detecting *dps* reporter signal alongside revTetR-mCherry. Because Dox inhibits translation, the amount of *dps* signal did not significantly increase with Dox treatment due to translational inhibition **(Figure 2C)**. However, we were able to conclude that slow-growing or growth-arrested cells had higher levels of revTetR mCherry signal using ATc and NO treatments.

We recently showed with a slow-folding fluorescent protein derivative (DsRed_42_) that NO stress was sufficient to promote fluorescence accumulation, but also found there was limited overlap in slow-growing and NO-stressed populations within host tissues (16). Destabilized and stable versions of fluorescent proteins were used to show that NO stress was transient within host tissues, and that few bacteria were actively responding and detoxifying NO at any given timepoint (16). However, the fact that Hmp^+^ bacteria preferentially survive doxycycline treatment suggests that, while NO exposure may be transient, there could also be sustained changes within cells exposed to high levels of NO (22). Although *Yersinia* repairs iron-sulfur cluster-containing proteins in response to this stress (51), there may be long-term damage, or even genetic changes that occur following NO exposure. It would be very interesting to determine if any genetic changes occur within microcolonies over the course of infection, especially given these bacterial populations are clonal, founded by a single bacterium, and presumed to be genetically homogenous (17).

## Materials and Methods

### Bacterial strains and growth conditions

*Y. pseudotuberculosis* IP2666 was used as the wild-type strain throughout. For *in vitro* experiments, overnight cultures were grown in LB at 26°C with rotation for 16 hours. Exponential phase cells for *in vitro* experiments were generated by sub-culturing overnight cultures by diluting 1:100 into fresh LB, and rotating at 37°C 2 hours. All *in vitro* experiments were performed by culturing at 37°C with rotation. For all mouse infections, bacterial inocula were grown overnight (16 hours) to post-exponential phase in 2xYT broth (LB, with 2x concentration of yeast extract and tryptone) at 26°C with rotation.

### revTetR reporter construction

The revTetR reporter was generated by introducing specific point mutations into our previously described *tetR::P_tetA_::mCherry* reporter construct in pMMB67EH (22). *tetR* was mutated by site-directed mutagenesis at two sites, E15A (GAG→GCU) and L17G (CUU→GGU), using 4 point mutations to generate the revTetR construct. An additional L25V mutation could not be successfully introduced. Primers were designed that contained the desired mutations, and were used to PCR amplify around the pMMB67EH plasmid using PfuUltra II high-fidelity polymerase (Agilent). PCR products were purified (QIAGEN) and transformed into electrocompetent DH5αλpir *E. coli.* Colonies were screened based on mCherry fluorescence in the absence of tetracyclines, plasmids were then isolated using the PureYield Plasmid Midiprep kit (Promega), and sequenced to confirm mutations. *revtetR::P_tetA_::mCherry* plasmids were transformed into *Y. pseudotuberculosis* strains as previously described (52), to generate the following reporter strains: *revtetR::P_tetA_::mCherry P_dps_::gfp-ssrA*, *revtetR::P_tetA_::mCherry gfp^+^*, and *revtetR::P_tetA_::mCherry P_hmp::_gfp*. *P_dps_::gfp-ssrA, gfp^+^*, and *P_hmp::_gfp* reporter constructs have been previously described (16), and are all inserted into the pACYC184 plasmid. *revtetR::P_tetA_::yfp* was constructed by PCR amplifying *revtetR::P_tetA_* and fusing this to *yfp* using overlap extension PCR. *revtetR::P_tetA_::yfp* was inserted into the pACYC184 plasmid, and transformed into a *Y. pseudotuberculosis* strain containing the previously described *P_hmp_::mCherry* plasmid (17).

### In vitro reporter characterization

Once subcultures reached exponential phase (2h growth), the indicated doses of doxycycline (Dox; 0.01 µg/mL, 0.1 µg/mL, or 1 µg/mL) or anhydrotetracycline (ATc; 0.1 µg/mL, 1 µg/mL, or 2 µg/mL) were added to samples to promote mCherry dilution, with one sample left untreated as a control. Samples were incubated at 37°C with rotation throughout the time course, and time points of 0, 0.5, 1, 2, 3, and 4h post-antibiotic treatment were taken. For each time point, 600 µL from each sample was pelleted and resuspended in 600 µL of PBS. 200 µL of each sample was then pipetted into 3 wells of a black walled, clear bottom 96 well plate, and analyzed using a Synergy Microplate Reader (Biotek Instruments) for OD_600_ and fluorescence signal measurements. Optical density (A_600nm_) was used to approximate cell number, and mCherry fluorescence was detected with 560nm excitation/610nm emission. All experiments were repeated in triplicate to generate a total of three independent data sets, or nine replicates. In strains of Yptb also containing a GFP reporter, GFP fluorescence was detected with 480nm excitation/520nm emission.

### Induction of hmp using nitric oxide donor compound

Overnight cultures were diluted 1:50 into fresh LB with the nitric oxide donor compound DETA-NONOate (2.5 mM dose) or were left untreated. After either 2 hours or 30 min of incubation, each culture was evenly split into two separate tubes. Samples were then further treated with either ATc (1 µg/mL dose) or left untreated, for a total of four different treatment groups: +NO/+ATc, +NO/-ATc, -NO/+ATc, - NO/-ATc. Aliquots were taken every 2h after ATc treatment to quantify absorbance and fluorescence, and cells were prepared for fluorescence microscopy, as described below. Fixed cells were imaged and quantified for single-cell fluorescence measurements.

### Fluorescence microscopy: bacteria

To visualize individual bacterial cells, 500 µL from each sample was pelleted, resuspended in 4% paraformaldehyde (PFA), and incubated overnight at 4° C for fixation. Agarose pads were prepared by placing two pieces of tape approximately 1 cm apart on slides, pipetting 25 µL of 1% agarose in PBS, and placing a glass coverslip over the area. After leaving the agarose to solidify for 20 min, the coverslips and tape were peeled off and the agarose was trimmed to 1 cm^2^ squares. PFA-fixed bacteria were pelleted, resuspended in 50 µL PBS, and 5 µL of sample was pipetted onto each agarose pad. Bacteria were imaged with the 63x oil immersion objective, using a Zeiss Axio Observer.Z1 (Zeiss) fluorescent microscope with Colibri.2 LED light source and an ORCA-R^2^ digital CCD camera (Hamamatsu). Five images were taken of each sample in distinct fields of view, using DIC, GFP, and mCherry channels. Volocity image analysis software was used to specifically select individual bacterial cells and quantify the fluorescent signal associated with each cell.

### Murine model of systemic infection

The following strains were used for mouse infections: *Y. pseudotuberculosis tetR::P_tetA_::mCherry*-*ssrA gfp^+^* (22)*, revtetR::P_tetA_::mCherry gfp^+^,* and *revtetR::P_tetA_::mCherry P_hmp::_gfp,* and *revtetR::P_tetA_::yfp P_hmp_*::mCherry. Six to eight week old female C57BL/6 mice were obtained from Jackson Laboratories (Bar Harbor, ME). All animal studies were approved by the Institutional Animal Care and Use Committee of Johns Hopkins University. Mice were injected intravenously with 10^3^ bacteria for all experiments. After 48 hours, mice were treated with the indicated dose (4mg/kg, equates to approximately 72µg in 100µl sterile PBS) of Dox or ATc, or left untreated. At the indicated timepoints, either 4h or 24h post-treatment, mice were euthanized via both lethal dose of isoflurane and cervical dislocation, and spleens were removed, halved, and processed for histology, or homogenized to quantify CFUs.

### Histology

Harvested spleen halves were fixed in 4% PFA overnight at 4° C, and were embedded using Sub Xero OCT compound (Mercedes Medical), frozen on dry ice, and stored at -80°C. Spleens were cross-sectioned into 10 um sections using a cryostat microtome; one representative section was imaged per mouse. To visualize reporters, sections were thawed in PBS, stained with Hoechst at a 1:10,000 dilution, washed in PBS, and coverslips were mounted using ProLong Gold (Life Technologies). Images were taken of all microcolonies within each section with a 63x oil immersion objective, using a Zeiss Axio Observer.Z1 (Zeiss) fluorescent microscope with Colibri.2 LED light source, an Apotome.2 (Zeiss) for optical sectioning, and an ORCA-R^2^ digital CCD camera (Hamamatsu).

### Image analysis

For *in vitro* experiments, Volocity image analysis software was used to select single bacterial cells in each image with the ‘find objects’ function. The mean and sum signal intensity was calculated for each object (single cell) for mCherry, GFP, and YFP channels. Sum signal intensity values were used for revTetR reporter quantification. For *in vivo* experiments, either Volocity or Fiji image analysis software was used to select regions of interest at the centroid and periphery of each microcolony and quantify the fluorescent signal intensity of each channel to generate relative signal intensities of fluorescent reporters. A small representative point (∼0.01pixels^2^) was selected at the centroid and periphery of each microcolony, which generated a ‘signal intensity’ value at that point that correlates with the sum. Centroid bacteria were defined as surrounded only by other bacteria. Peripheral bacteria were defined based on contact with host cells (**Figure 4D**). All data was analyzed and graphed, and all statistical analyses performed, using GraphPad Prism 9.

## Author Contributions

Conceptualization: BL, RKD, KMD; Formal Analysis: BL, REB, KMD; Funding Acquisition and Supervision: KMD; Investigation: BL, REB, KLC, KMD; Methodology: BL, REB, KMD; Writing – Original Draft Preparation: BL, REB, KMD; Writing – Review & Editing: BL, REB, KLC, RKD, KMD.

## Acknowledgments

We thank the members of the Davis lab, who provided feedback and suggestions during the final steps of manuscript preparation. We also thank the members of the Pekosz and Klein lab for their helpful feedback throughout this project. This work was supported by NIAID grants 1K22AI123465-01 and 1R21AI154116-01A1 to KMD. REB is also supported by training grant 2T32AI007417-26 through NIAID. The authors of this manuscript declare no conflicts of interest.

