## Supplemental Materials for "NO-stressed *Y. pseudotuberculosis* have decreased cell division rates in the mouse spleen"

### Supplemental Material

#### Methods associated with Supplemental Figures

*Induction of hmp using nitric oxide donor compound.* Overnight cultures were diluted 1:50 into fresh LB with the nitric oxide donor compound DETA-NONOate (2.5 mM dose) or were left untreated. After either 2 hours or 30 min of incubation, each culture was evenly split into two separate tubes. Samples were then further treated with either ATc (1 μg/mL dose) or left untreated, for a total of four different treatment groups: +NO/+ATc, +NO/-ATc, -NO/+ATc, -NO/-ATc. Aliquots were taken every 2h after ATc treatment to quantify absorbance and fluorescence, and cells were prepared for fluorescence microscopy, as described below. Fixed cells were imaged and quantified for single-cell fluorescence measurements.

#### Supplemental Figure Legends

**Supplemental Figure 1. Hmp reporter signal is heightened in bacteria treated with the NO donor compound.** Cultures of the revTetR-mCherry $P_{hmp}$*::gfp* strain were cultured two hours in the presence of NO (+NO treatment), then split, and either treated with 1μg/ml ATc (+ATc) or left untreated (-ATc). Aliquots of cells were fixed at the indicated timepoints after ATc addition for fluorescence microscopy. Single cell fold change in GFP signal with NO treatment is shown, compared to untreated cells. Dotted line at a value of 1 is the mean GFP signal from individual untreated cells (baseline signal). Median value/group is highlighted.

**Supplemental Figure 2. The revTetR-YFP signal no longer dilutes when cells reach stationary phase.** Overnight cultures of the revTetR-YFP $P_{hmp}$*::mCherry* strain were diluted 1:50 and grown in the presence (+NO) or absence (-NO) of NO for 2h. Cultures were then further split with 1 µg/ml ATc (+ATc) or left untreated, and aliquots taken at each indicated timepoint after ATc addition. **(A)** Sum single cell YFP signals and the corresponding **(B)** mean single cell mCherry signals are shown with the mean signals highlighted, as determined by microscopy image analyses (Volocity). At least 1,300 individual cells were quantified per group. **(A-B)** Kruskal-Wallis one-way ANOVA with Dunn's post-test was used to evaluate the significance of differences between groups; ****p<.0001.

**Supplemental Figure 3. RevTetR-YFP signal accumulates in both slow-growing and fast-growing cells in the absence of ATc.** Overnight cultures of the revTetR-YFP $P_{hmp}$*::mCherry* strain were grown in the presence (+NO) or absence (-NO) of NO for 30 min. Cultures were then further split with 1 µg/ml ATc (+ATc) or left untreated, and aliquots taken at each indicated timepoint after ATc addition. **(A)** Sum single cell YFP signal is shown along with the corresponding **(B)** mean single cell mCherry signal (Hmp reporter) with the mean signals highlighted, as determined by microscopy image analyses (Volocity). At least 1,500 individual cells were quantified per group. **(A-B)** Kruskal-Wallis one-way ANOVA with Dunn's post-test was used to evaluate the significance of differences between groups; ****p<.0001.

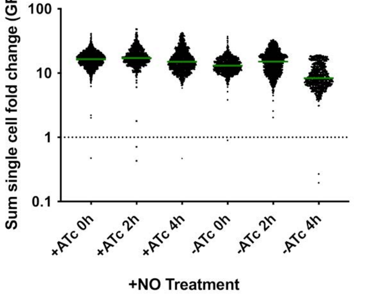

100
10
1
0.1
Sum single cell fold change (GFP)
+ATc 0h
+ATc 2h
+ATc 4h
-ATc 0h
-ATc 2h
-ATc 4h
+NO Treatment

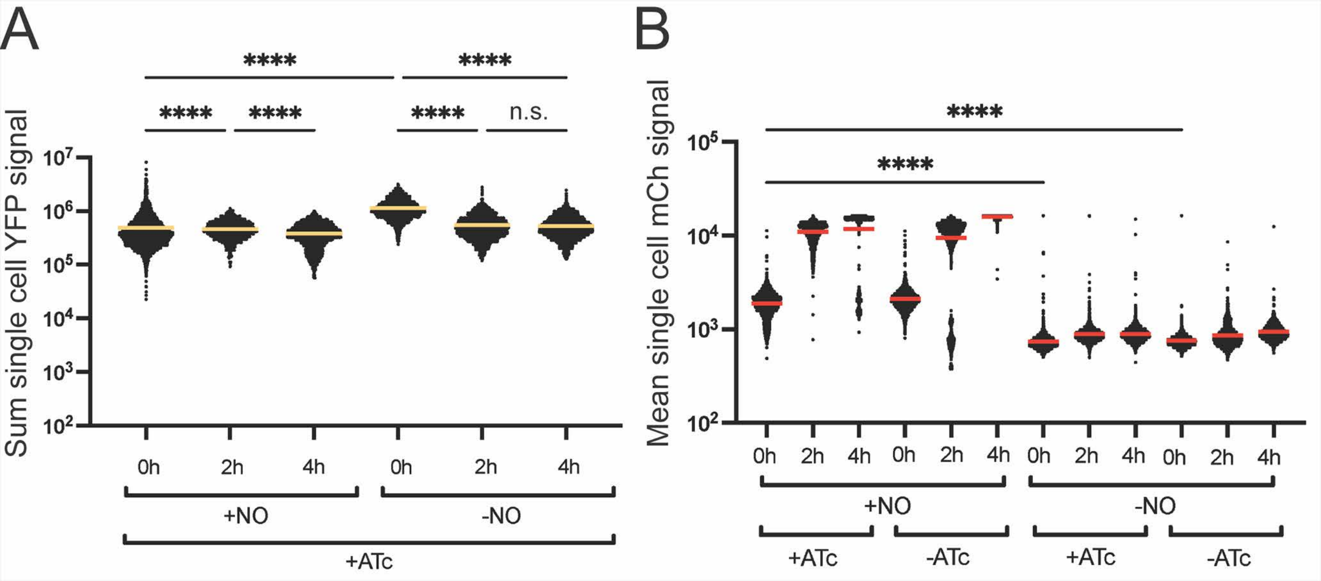
A
B
****
****
****
****
****
n.s.
Sum single cell YFP signal
$10^7$
$10^6$
$10^5$
$10^4$
$10^3$
$10^2$
0h
2h
4h
0h
2h
4h
+NO
-NO
+ATc
****
****
Mean single cell mCh signal
$10^5$
$10^4$
$10^3$
$10^2$
0h
2h
4h
0h
2h
4h
0h
2h
4h
0h
2h
4h
+NO
-NO
+ATc
-ATc
+ATc
-ATc

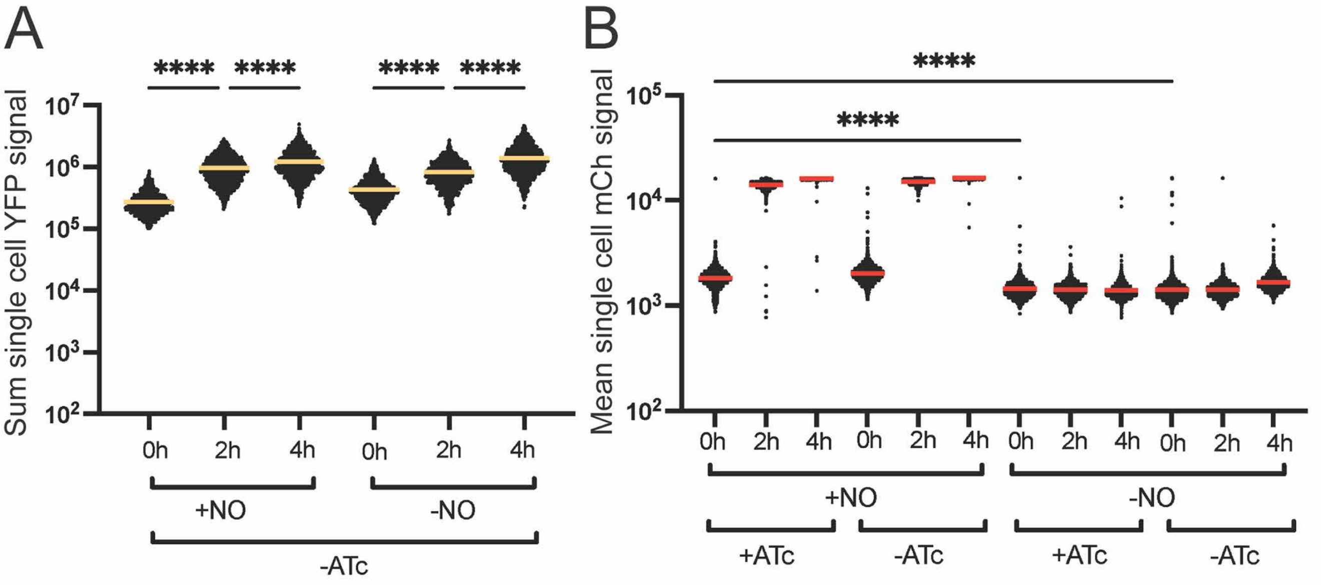
A
****
****
****
****
Sum single cell YFP signal
$10^7$
$10^6$
$10^5$
$10^4$
$10^3$
$10^2$
0h
2h
4h
0h
2h
4h
+NO
-NO
-ATc
B
****
****
Mean single cell mCh signal
$10^5$
$10^4$
$10^3$
$10^2$
0h
2h
4h
0h
2h
4h
0h
2h
4h
0h
2h
4h
+NO
-NO
+ATc
-ATc
+ATc
-ATc
